# Targeted long-read sequencing facilitates phased diploid assembly and genotyping of the human T cell receptor alpha, delta and beta loci

**DOI:** 10.1101/2022.05.24.493244

**Authors:** Oscar L. Rodriguez, Catherine A. Silver, Kaitlyn Shields, Melissa L. Smith, Corey T. Watson

## Abstract

T cell receptors (TCRs) recognize peptide fragments presented by the major histocompatibility complex (MHC) and are critical to T cell mediated immunity. Early studies demonstrated an enrichment of polymorphisms within TCR-encoding (TR) gene loci. However, more recent data indicate that variation in these loci are underexplored, limiting understanding of the impact of TR polymorphism on TCR function in disease, even though: (i) TCR repertoire signatures are heritable and (ii) associate with disease phenotypes. TR variant discovery and curation has been difficult using standard high-throughput methods. To address this, we expanded our published targeted long-read sequencing approach to generate highly accurate haplotype resolved assemblies of the human TR beta (TRB) and alpha/delta (TRA/D) loci, facilitating the detection and genotyping of single nucleotide polymorphisms (SNPs), insertion-deletions (indels), structural variants (SVs) and TR genes. We validate our approach using two mother-father-child trios and 5 unrelated donors representing multiple populations. Comparisons of long-read derived variants to short-read datasets revealed improved genotyping accuracy, and TR gene annotation led to the discovery of 79 previously undocumented V, D, and J alleles. This demonstrates the utility of this framework to resolve the TR loci, and ultimately our understanding of TCR function in disease.

## Introduction

T cell receptors (TCR) play a central role in the adaptive immune system and are critical for fighting pathogens(1). TCRs are expressed on the T cell surface and interact with antigens via major histocompatibility complex (MHC) proteins. TCRs exist as a heterodimer protein consisting of either paired alpha and beta chains, or paired gamma and delta chains. The total number of genes that encode the human TCR chains ranges from 228 to 234 and are grouped into four segment types: variable (V), diversity (D), joining (J) and constant (C) genes. These genes reside in three different genomic loci(2). During T cell development, a V, D and J gene are selected during V(D)J recombination from the beta (TRB) and delta loci (TRD), and a V and J gene from the alpha (TRA) and gamma (TRG) loci; the recombined genes serve as the template for the transcription and translation of a given TCR. The immense diversity observed in the TCR repertoire (i.e. the complete set of TCRs) is seeded by the selection from a large number of V, D and J genes within the TCR loci, combined with junctional diversity at V-D and D-J junctions, and allows T cells to mount an immune response to diverse antigens(3). It is estimated that 1×10^13^ unique TCRs exist in a single individual(4). Development of the overall TCR repertoire is shaped by host genetics and the environment, including foreign and self-peptides(5–7).

The extent to which genetics affects the TCR repertoire and the downstream impact of this genetic diversity on the ability to mount effective disease-related T cell responses remains unknown. Several targeted genetic studies have identified germline variants associated with TCR function and TCR repertoire features. For example, TCRs using the allele TRBV9*02 have been shown to have reduced functional recognition of Epstein-Barr virus as compared to TRBV9*01 under the same human leukocyte antigen (HLA) background(8). Furthermore, single nucleotide polymorphisms (SNPs) within TCR and HLA have been associated with differences in TCR V gene usage(6, 9), and TCR repertoires in monozygotic twins have been shown to be more similar than repertoires of unrelated individuals(5), demonstrating a role for genetics in the development of the TCR repertoire. However, large scale genome-wide association studies (GWAS) have only implicated the TCR loci twice, specifically in narcolepsy(10, 11) and renal function after transplantation(12).

Several reasons could exist for the disconnect between TCR polymorphisms and phenotypic outcomes, including but not limited to, small sample sizes, inadequate genotyping, the use of an incomplete reference assembly, and disease/sample heterogeneity. Incomplete genotyping could be due to the complex repetitive nature of the antigen receptor loci. Similar to the immunoglobulin (IG) loci, the TCR loci have been shaped by gene duplication events(13, 14), which have resulted in large expansions of the TCR gene family. Due to the duplicated and repetitive structure of immune receptor loci such as TCR and IG, it has been proposed that nextgeneration short read sequencing (NGS) performs suboptimally in these regions(15, 16). Specifically in the IG heavy chain locus (IGH), we have previously demonstrated that the use of NGS results in high rates of false positive and negative variants(17). Because of these technical barriers we have not characterized the full extent of genetic diversity in these immune receptor loci(9), limiting our understanding of the role of germline variation in TCR function.

Here, we describe the application of a newly developed probe-based capture design to conduct long-fragment targeted enrichment of the TRB and TRA/D loci for high fidelity long read single molecule real time (SMRT) sequencing. To provide a proof-of-concept dataset demonstrating the unique value of this approach, we have generated targeted SMRT sequencing data on two trios and five unrelated individuals of African, Asian and European ancestry. First, we show the efficacy of this capture-based sequencing approach and how it can be used to generate haplotype-resolved assemblies. We then use these assemblies to identify variants of different classes, including SNPs, indels, and structural variants (SVs), and resolve and curate full length genes and alleles. Using variants resolved from short-read NGS from the same individuals, we compare the concordance between sequencing methods. This study demonstrates a robust and accurate methodology to efficiently resolve the TRB and TRA/D loci, validates previously identified variants, and takes a step forward in providing a framework that will be effective at resolving the role of TRB and TRA/D genetics in TCR-related phenotypes.

## Methods

### Long-read library preparation and sequencing

Genomic DNA was procured from the Coriell Institute for Medical Research (Camden, NJ). Genomic DNA was prepared using the protocol described in our previously published targeted long-read sequencing and IGenotyper framework(17). Briefly, 1-2 micrograms of high molecular weight DNA was sheared using g-tube (Covaris, Woburn, MA) to 5-9 Kbp at a 7000 RPM and size selected using the 0.75% DF 3-10kb Marker S1-Improved Recovery cassette definition on the Blue Pippin (Sage Science). Sheared gDNA underwent end repaired and A-tailing using the standard KAPA library protocol. Barcodes were added to samples sequenced on the Sequel or SequelII platform and universal primers were ligated to all samples. PCR amplification was performed for 8-9 cycles using PrimeSTAR GXL Polymerase (Takara) at an annealing temperature of 60°C. Small fragments and excess reagents were removed using 0.7X vol:vol AMPure beads. Libraries were captured and washed using the KAPA HyperCap protocol, and post-capture PCR amplification was performed for 16-18 cycles using PrimeSTAR GXL Polymerase (Takara) at an annealing temperature of 60°C.

Capture libraries were prepped using SMRTbell Template Preparation Kit 1.0 (Pacific Biosciences, Menlo Park, CA, United States). Each sample was treated with a DNA Damage Repair and End Repair mix to repair nicked DNA, followed by A-tailing and ligation with SMRTbell hairpin adapters. These libraries were treated with an exonuclease cocktail to remove unligated gDNA and cleaned with 0.5X AMPure PB beads (Pacific Biosciences). The resulting SMRTbell libraries were prepared for sequencing according to the manufacturer’s protocol and sequenced on the Sequel lie system using 2.0 chemistry and 30 h movies. HiFi data, consisting of circular consensus sequences filtered at a quality threshold of QV20 (99%), were generated on instrument and used for all downstream analysis.

### Long-read assembly and genetic variation and allele detection

HiFi sequence data was analyzed using IGenotyper(17). IGenotyper uses BLASR, WhatsHap, MsPAC and Canu for alignment, SNP detection and phasing, HiFi phasing and indel/SV detection and assembly, respectively(41–44). Briefly, the HiFi reads were first aligned to GRCh38 containing the alternate contigs. WhatsHap was then used to detect phased SNPs using the find_snv_candidates, genotype and phase commands. Using the MsPAC phase scripts, the HiFi reads were partitioned by haplotype or were labeled as unphased using the BLASR aligned reads and the WhatsHap phased SNPs. Canu was then used to assemble each haplotype and unphased reads with the ‘-pacbio-hifi’ option. MsPAC was then used again to detect indels and SVs using a multiple sequence alignment generated by Kalign(45). SNPs were detected by IGenotyper from the alignment of haplotype resolved assemblies.

TR genes and alleles were annotated and genotyped using IGenotyper. Specifically, IGenotyper output included extracted allele sequences for each TR gene from the assembly and HiFi reads; allele assignments for each gene were made by assessing whether extracted sequences were exact string matches to the IMGT database. The IMGT database used was downloaded on 2022-02-20. Sequences that did align to IMGT with an exact match were considered novel. Novel alleles were also compared to the pmIGTR database using the fasta sequences found in https://pmtrig.lumc.nl/Analytics, again by requiring exact sequence matches.

### Assessing accuracy of assembly and genetic variants

The accuracy of the assemblies and variants was assessed using data from two mother-father-proband trios. In each case, we used BLASR(41) to align the inherited parental contigs to the respective maternally or paternally inherited assembly in the proband(41). Differences between the parental and proband sequences were detected using a custom python script utilizing the pysam library. Alignments were evaluated for (1) full alignment without any soft-clipped sequences, (2) deletions, (3) insertions and (4) mismatches.

### Comparison of SNP genotypes between long- and short-read datasets

1KGP3 and 1KG-30X SNPs were downloaded from s3://1000genomes/release/20130502/supporting/GRCh38_positions/ALL.chr7.phase3_shapeit2_mvncall_integr ated_v3plus_nounphased.rsID.genotypes.GRCh38_dbSNP_no_SVs.vcf.gz and gs://fc-56ac46ea-efc4-4683-b6d5-6d95bed41c5e/CCDG_14151/Project_CCDG_14151_B01_GRM_WGS.gVCF.2020-02-12/Project_CCDG_14151_B01_GRM_WGS.gVCF.2020-02-12, respectively. Comparisons were performed using a custom python script that compared both the presence and genotype of all SNPs between long-read and short-read datasets.

## Results

### Targeted long-read sequencing of the TRA/D and TRB loci

We designed a custom oligo capture panel (Roche KAPA HyperChoice) based on sequence targets spanning the TRA/D and TRB loci (hg38, chr22:22000934-22953034; chr7:142270924-142843399). To demonstrate the utility of this approach, we generated long-fragment capture libraries (4-6 Kb) in 11 individuals from the 1000 Genomes Project (1KGP) cohort, including 2 mother-father-child trios (Yoruba, n=3; Kinh Vietnamese, n=3) and 5 unrelated individuals of diverse ethnic backgrounds (Table 1; Japanese, n=1; Yoruba, n=1; European, n=1; Columbian, n=1; Puerto Rican, n=1). Libraries were multiplexed and sequenced on the Pacific Biosciences Sequel IIe system, generating mean high fidelity (HiFi) read lengths and accuracy ranging from 4.2 to 6.0 Kb (Figure 1a) and 99.90 to 99.93% (Figure 1b), respectively. Average per base HiFi read coverage ranged from 9X to 246X across both loci (Table 1). Only 227 bases collectively across TRA/D and TRB (0.014%) were found to have 0X coverage in just two of the samples sequenced; in the remaining nine samples, all bases were spanned by at least 1 HiFi read (Figure 1c). In summary, these metrics demonstrate that TRA/D and TRB are effectively captured and sequenced using this targeted custom oligo capture panel and long-read sequencing framework.

**Figure 1.**
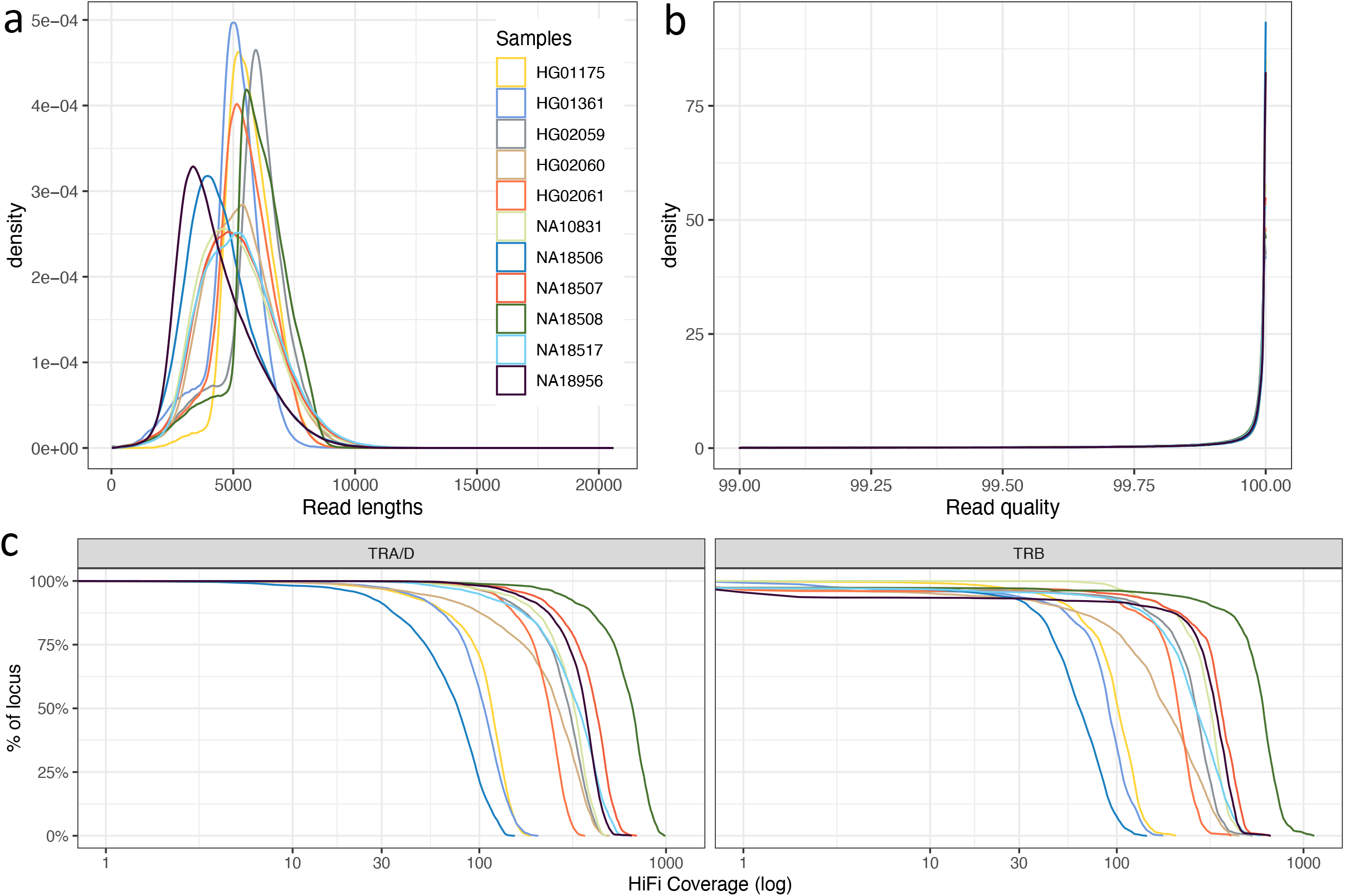
Targeted HiFi sequencing generates highly accurate long reads. HiFi read (**A**) lengths and **(B)** quality. Coverage of (**C**) TRA/D and (**D**) TRB. Each color represents a different sample.

**Table 1.**
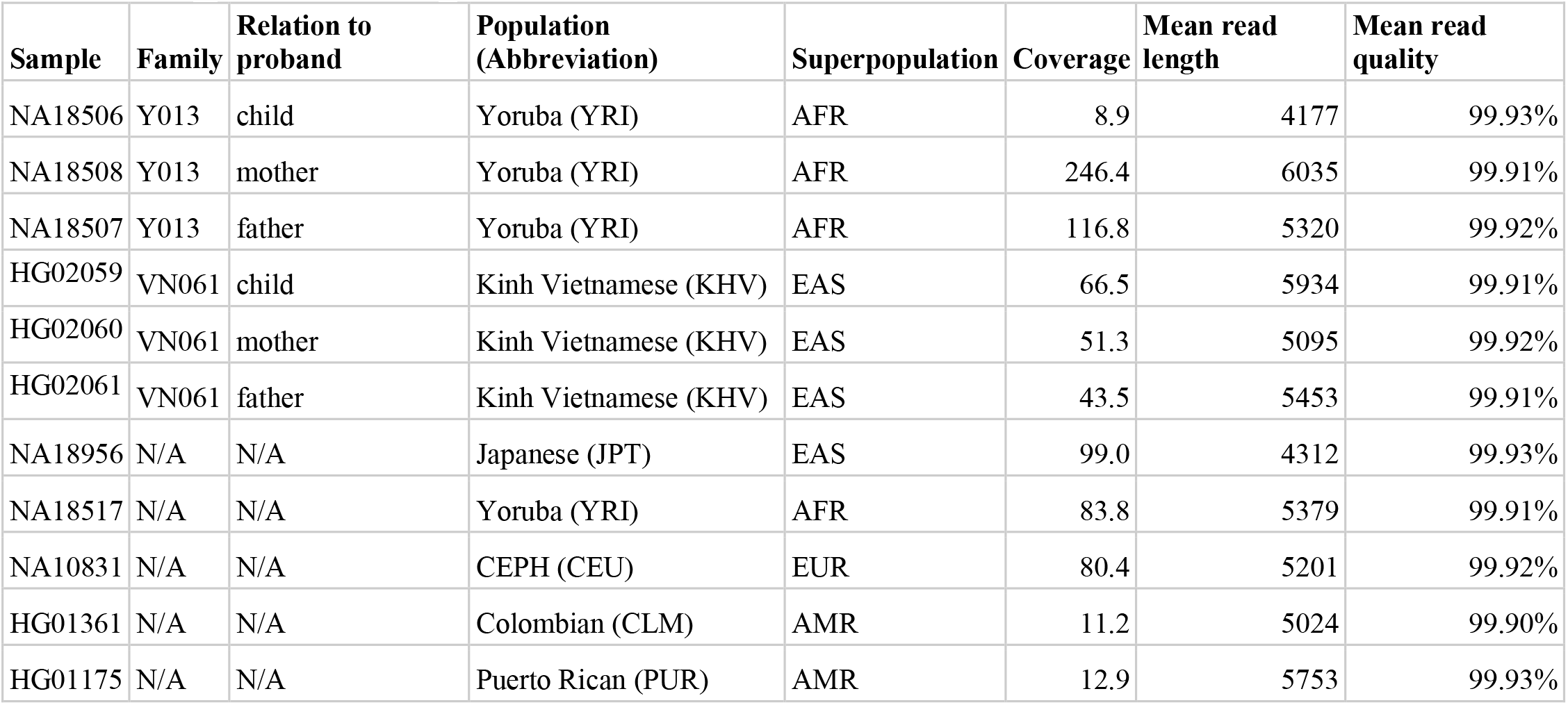
Samples used in study.

### Generating haplotype-resolved assemblies of the TRA/D and TRB loci

We next assessed our ability to generate TRA/D and TRB haplotype-specific assemblies. Following methods outlined in Rodriguez et al. 2020, heterozygous SNPs were identified and phased using HiFi reads from the capture-based sequencing datasets. Among the unrelated samples (n=9), a mean of 1314 (TRA/D) and 1110 (TRB) heterozygous SNPs were identified. On average, >99% of these SNPs (1353, TRA/D;1105, TRB) were phased with the WhatsHap tool allowing for 57 to 93% (mean=78%) and 29 to 95% (mean=66%) of TRA/D and TRB, respectively, to be completely phased. The variation in assembly phasing was associated with the number of heterozygous positions identified in each sample. For example, NA18956, a Japanese sample, had the fewest bases phased (29% in TRB), and also had the fewest heterozygous SNPs (n=463). However, the low number of polymorphisms identified was not due to low read coverage as this sample had 99X mean base coverage. The longest haplotype blocks resolved with this approach represented 71.0% and 83.0% of the TRA/D and TRB loci, respectively, revealing that, for at least some samples, the majority of these loci could be phased into a single block. The largest assembled contigs per sample ranged in size from 158 to 653 Kb (mean = 295 Kb) and from 100 Kb to 326 Kb (mean = 180 Kb) in TRA/D and TRB, respectively, showing that, in some samples, large portions of haplotypes spanning these loci were assembled into a single contig.

We next evaluated whether the trio probands HG02059 and NA18507 could be fully phased using SNPs phased with paternal and maternal data. In both probands, both haplotypes in TRA/D and TRB were completely phased and resolved, including 20.2 Kb and 20.4 Kb insertions found in both proband haplotypes in TRB. The accuracy of the TRA/D and TRB assemblies were assessed using parental contigs (Figure 2). The HG02059 maternally-inherited haplotypes only had 63 base pair mismatches with the HG02060 (mother) contigs, collectively across the TRA/D and TRB loci, representing an assembly accuracy greater than 99.996% (1491720/1491783 bp). Likewise the paternally-inherited haplotypes for TRA/D and TRB only had 55 base pair mismatches with HG02061 (father) contigs, representing an assembly accuracy of 99.997% (1642320/1642375). Results for assemblies generated from NA18507 were similar, with 99.996% accuracy for both loci and maternal/paternal haplotypes. Together, these results indicate that TRA/D and TRB can be completely reconstructed at high quality using our approach.

**Figure 2.**
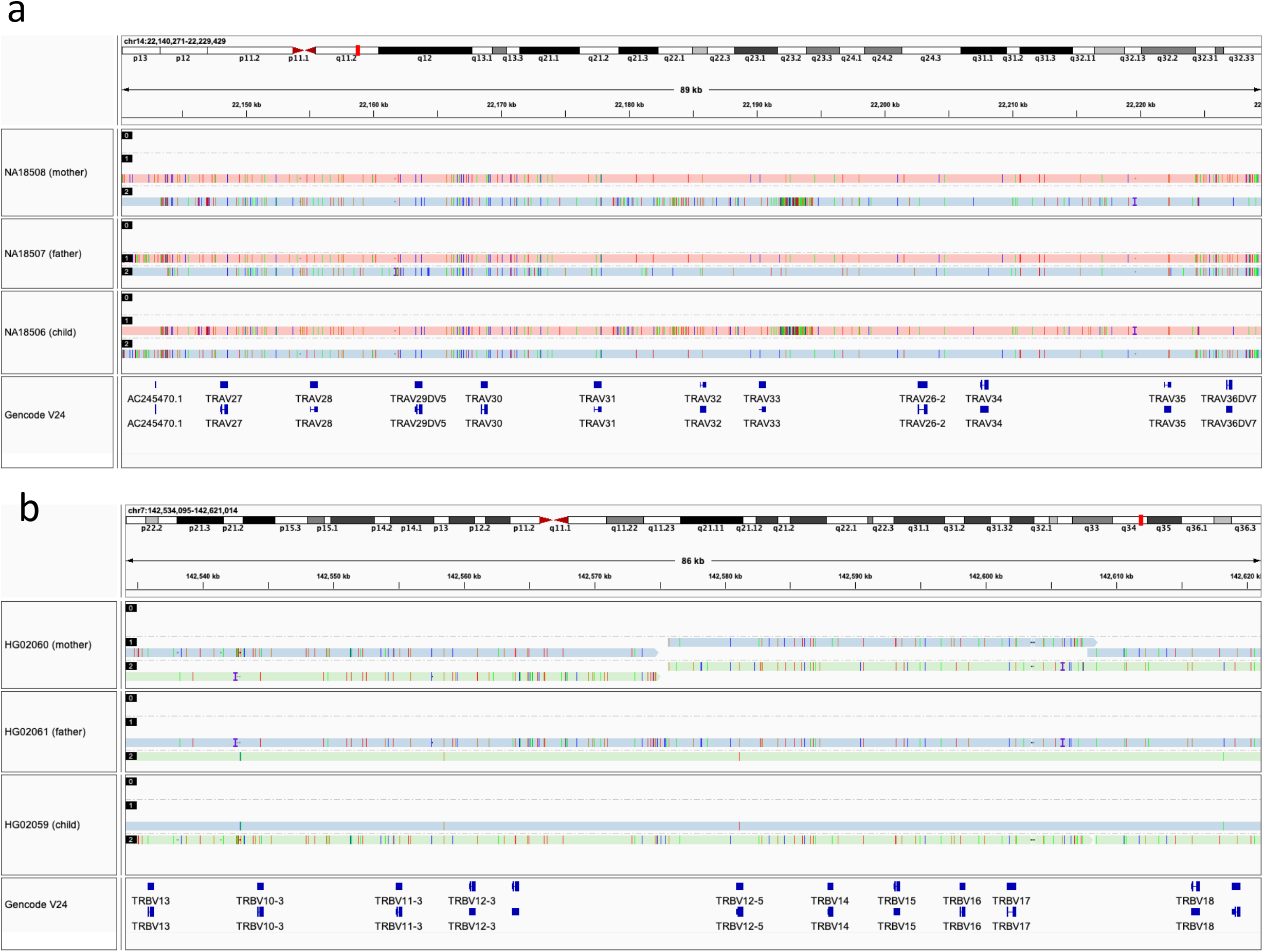
Trio assemblies in TRA/D and TRB are highly accurate. Examples of high concordance in TRA/D (**A**) and TRB (**B**) assemblies are shown between the parents and proband of both trios.

### Accurate detection of SNPs, indels and SVs from diploid assemblies

Using the haplotype-specific assemblies, genetic variants including SNPs, indels (2-49 bp), and SVs (>= 50 bp) were evaluated. First the accuracy of the genetic variants was assessed using Mendelian inheritance in the two trios. There were 4065 (99.50%) and 4571 (99.13%) SNPs that followed proper Mendelian inheritance in HG02059 and NA18506, respectively. In contrast, only 22 and 40 SNPs did not follow proper Mendelian inheritance. For indels, the Mendelian inheritance rate was 97% (382/394) and 94% (448/475) in HG02059 and NA18506, respectively, and for SVs it was 93% (14/15) and 95% (19/20). These data show that variants within TRA/D and TRB are being accurately detected.

Across the unrelated samples (n=9), the mean numbers of SNPs, indels and SVs identified per sample in TRA/D and TRB, respectively, were 2470 and 1802, 260 and 132, and 4.2 and 6.3 (Figure 3a-c). As expected, the majority of SNPs and indels resided within intergenic regions, but in each sample we observed a mean of 36.33 and 61.44 SNPs within TRA/D and TRB genes. The total number of nonredundant SNPs in TRA/D and TRB was 5072 and 3385, respectively, of which only 10 and 16 SNPs were not found in dbSNPv154. However, 818 (16%) and 1206 (36%) SNPs in TRA/D and TRB, respectively, do not contain any allele frequency data. This indicates that while previous studies using large cohorts have identified these SNPs, they have not resolved them in enough samples to determine their allele frequency. Further, these SNPs are likely not rare, as 1443 (71%) were found in 2 or more of the 9 unrelated samples in this study.

**Figure 3.**
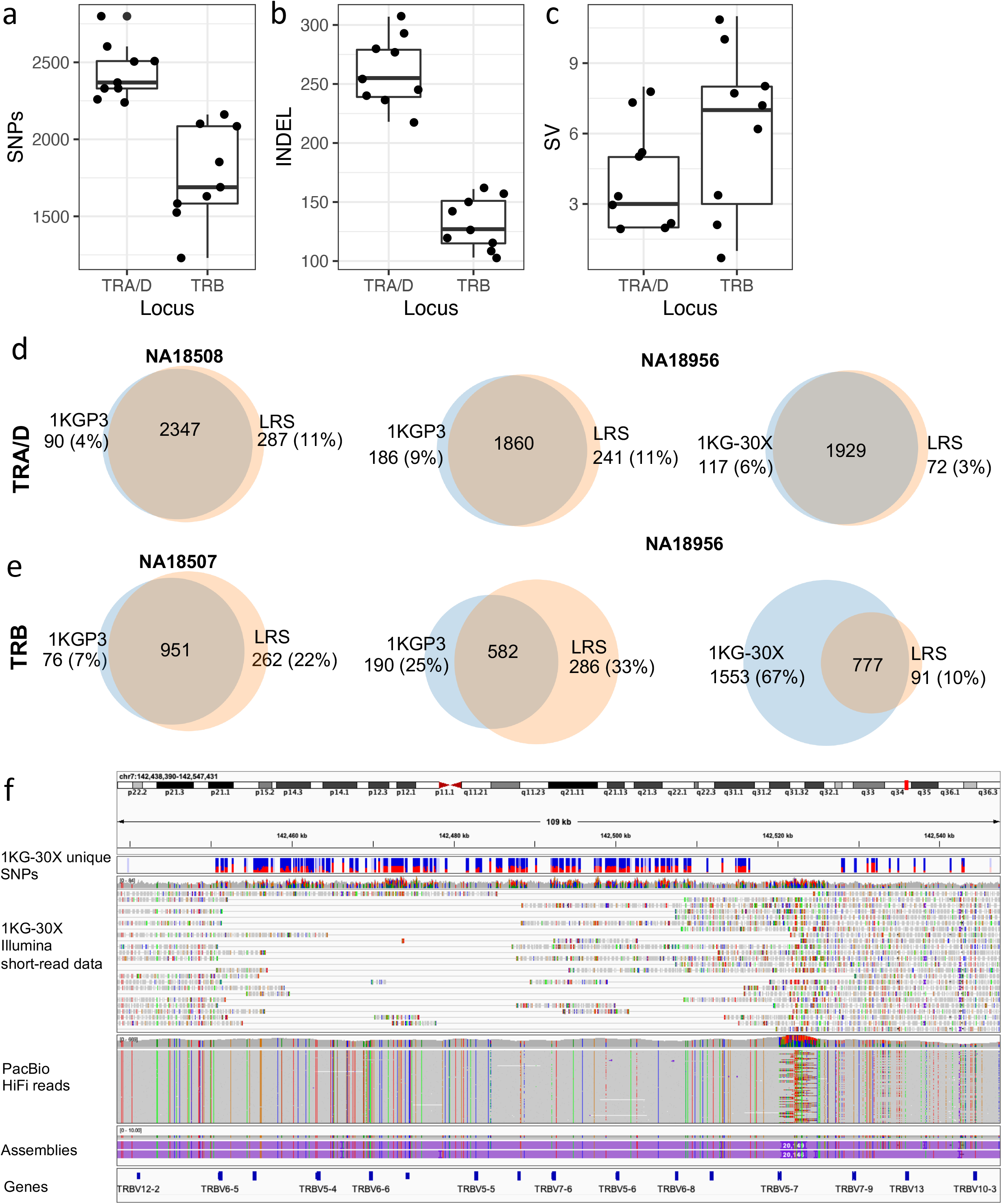
Genetic variants in TRA/D and TRB. The per sample counts of (**A**) SNPs, (**B**) indels and (**C**) SVs detected from TRA/D and TRB assemblies. Comparison of SNPs detected from long-read capture assemblies and 1KGP3 (left) and 1KG-30X (right) variant call sets for (**D**) TRA/D and (**E**) TRB; samples with the highest (left) and lowest (right) concordance rates are shown. (**F**) IGV screenshot showing example region of TRB (coords), demonstrating the presence of false SNPs identified from the 1KG-30X dataset. The panels shown are (1, top) SNPs present only within the 1KG-30X variant call set, (2) 1KG-30X Illumina 30X 2×151 bp shortread data, (3) PacBio HiFi-reads and (4, bottom) PacBio HiFi assemblies.

We also noted that SNP density in TRA/D and TRB was higher than what was observed across the entirety of chromosome 14 and 7. For 6 samples, we used the 1000 Genomes Project Phase 3 (1KP3) SNPs, to calculate the number of SNPs in 10 KB windows across chromosomes 14 and 7, representing a background SNP density for comparisons to the TRA/D and TRB loci, respectively. The mean number of SNPs per 10 Kb window across samples for chromosomes 14 and 7 ranged from 15 to 18 for both chromosomes. In TRA/D and TRB, SNP densities ranged from 25 to 30 and 20 to 26, respectively (Supplementary Figure 1). Interestingly, the increased SNP density was not uniformly distributed across the loci. Both TRA/D and TRB contained regions with an elevated number of SNPs (Supplementary Figure 1). For example, all samples contained 71 to 87 SNPs in the 10KB window spanning chr14:21890001-21900000 (TRA/D), which contains the gene *TRAV8-4*. Similarly, the 10KB window spanning chr7:142380001-142390000 in TRB, including the genes *TRBV6-4, TRBV7-3* and *TRBV5-3*, contained 23 to 83 SNPs across all unrelated samples. Furthermore, although SNP densities in TRA/D and TRB were elevated, runs of homozygosity (ROH) were also observed in both loci, with the longest ROHs observed in TRB (Supplementary Figure 2,3). This could be related to previously reported differences in recombination rates between TRA/D and TRB(18) (2.3 cM/Mb vs. 0.3 cM/Mb), as genomic regions with lower than average recombination rates have been shown to have longer tracts of ROH(19).

We identified three SVs >20 Kb in the TRB locus, two of which were polymorphic among the 9 unrelated samples. One of these polymorphic SVs involved the deletion of the genes *TRBV6-2, TRBV3-2*, and *TRBV4-3*. We also detected a number of smaller non-gene-containing intergenic SVs, the largest of which was a 1044 bp deletion. The largest intergenic insertion was 665 bps, represented by a tandem repeat expansion of a 37 bp motif. Other SVs included insertions and deletions of mobile element sequences (ME). For example, a 331 bp Alu insertion was found in 6 individuals. Additional SVs in non-repetitive regions were also found in multiple samples. For example, an 800 bp deletion and 592 bp insertion were found in 5 and 6 individuals, respectively. The positions of these SVs and their genotypes across samples are provided in Supplementary Table 4.

### Comparison between long-read and short-read derived variants

We have previously reported a high number of false-positive and -negative SNPs from short-read genomic data in the IGH locus(17). Given that the IGH and TCR loci are evolutionary related, sharing similar structural characteristics with respect to repeat sequences and segmental duplications, we wanted to determine whether SNPs identified with short-read data within the TCR loci might also be impacted by higher falsepositive and -negative rates. To assess this, we compared SNP genotype calls derived from our long-read capture data in 6 unrelated individuals to those called from two independent short-read datasets in these same samples: (1) Phase 3 variant call sets from the 1KGP (referred to as 1KGP3)(20), and (2) variants called from more recently generated 30X WGS Illumina NovaSeq 6000 2 x 151 paired-end, TruSeq PCR-free sequence data (referred to as 1KG-30X; Figure 3d,e)(21).

The percentage of 1KGP3 TRA/D and TRB SNPs present in the long-read SNP dataset ranged from 94 to 96% and 84 to 93%, respectively (Figure 3d,e). The number of SNPs identified solely by the long-read capture data when compared to the 1KGP3 call set ranged from 268 to 301 (11-14%) and 228 to 385 (22-36%) in TRA/D and TRB, respectively (Figure 3d,e). This shows that the 1KGP3 call set suffers from high falsepositive and -negative rates in TRB, and a high false-negative rate in TRA/D.

For the 1KG-30X call set, the number of SNPs detected solely by long-reads decreased substantially for both loci. For TRA/D and TRB, the SNPs identified solely by long-reads ranged from 80 to 105 (3.6 - 4.3%) and 12 to 39 (1.3 - 3.8%) respectively (Figure 3d,e). In addition, the number of 1KG-30X SNPs also identified by the long-read SNP datasets ranged from 97 to 99% (Figure 3d,e). However, for TRB, the opposite was true; the percentage of 1KG-30X SNPs found in the long-read SNP datasets ranged from 25 to 34% (Figure 3d,e). We determined that the significant increase in false-positive SNPs in 1KG-30X was related to the reference genome used. For the 1KGP3 call set, GRCh37/hg19 was used, and for the 1KG-30X call set, GRCh38/hg38 was used. GRCh37/hg19 has three ~20 Kb insertions in TRB relative to GRCh38/hg38. We observed that, in the 1KGP3-30X data set, reads derived from the ~20 Kb insertions were misaligned to other regions of TRB in the GRCh38/hg38 reference (Figure 3f); this was despite the fact that an alternate TRB haplotype with the insertion sequences was present (chr7_KI270803v1_alt) in the reference file used to generate the 1KG-30X call set.

### Large structural differences between three TRB reference haplotypes

Large structural differences between GRCh38/hg38 and GRCh37/hg19 references within the TRB regions have been noted previously(22, 23). In the previous section, we demonstrated that a significant number of false-positive SNPs from short read data localized around large ~20 Kb insertions absent from the primary chromosomes in GRCh38/hg38. We thus conducted a focused analysis of these regions here to more directly assess their support in the capture assemblies and their impacts on short-read mapping and variant detection. Between GRCh38/hg38 and GRCh37/hg19 reference files, there are 3 haplotypes available for TRB. Comparing the GRCh37/hg19 and chr7_KI270803v1_alt alternate contig to GRCh38/hg38 revealed an insertion and gaps in GRCh37/hg19 and 3 ~20 Kb insertions in chr7_KI270803v1_alt (Figure 4a,b). We assessed whether these events were supported by the HiFi read data generated here, by mapping these to the chr7_KI270803v1_alt haplotype (Figure 4c). We found that no inversions were detected, suggesting that this event is either a rare SV or a mis-assembly. False inversions have also been identified in other previously characterized misassembled regions in the human genome reference (24, 25). The chr7_KI270803v1_alt TRB haplotype additionally appears to be a minor haplotype as only 2 out 18 haplotypes have all three ~20 Kb insertions. However, it will be beneficial to use the chr7_KI270803v1_alt sequence, as it represents the longest haplotype and contains 5 functional genes (*TRBV6-2, TRBV4-3, TRBV6-9, TRBV7-8* and *TRBV5-8*) and one pseudogene (*TRBV3-2*; Figure 4d).

**Figure 4.**
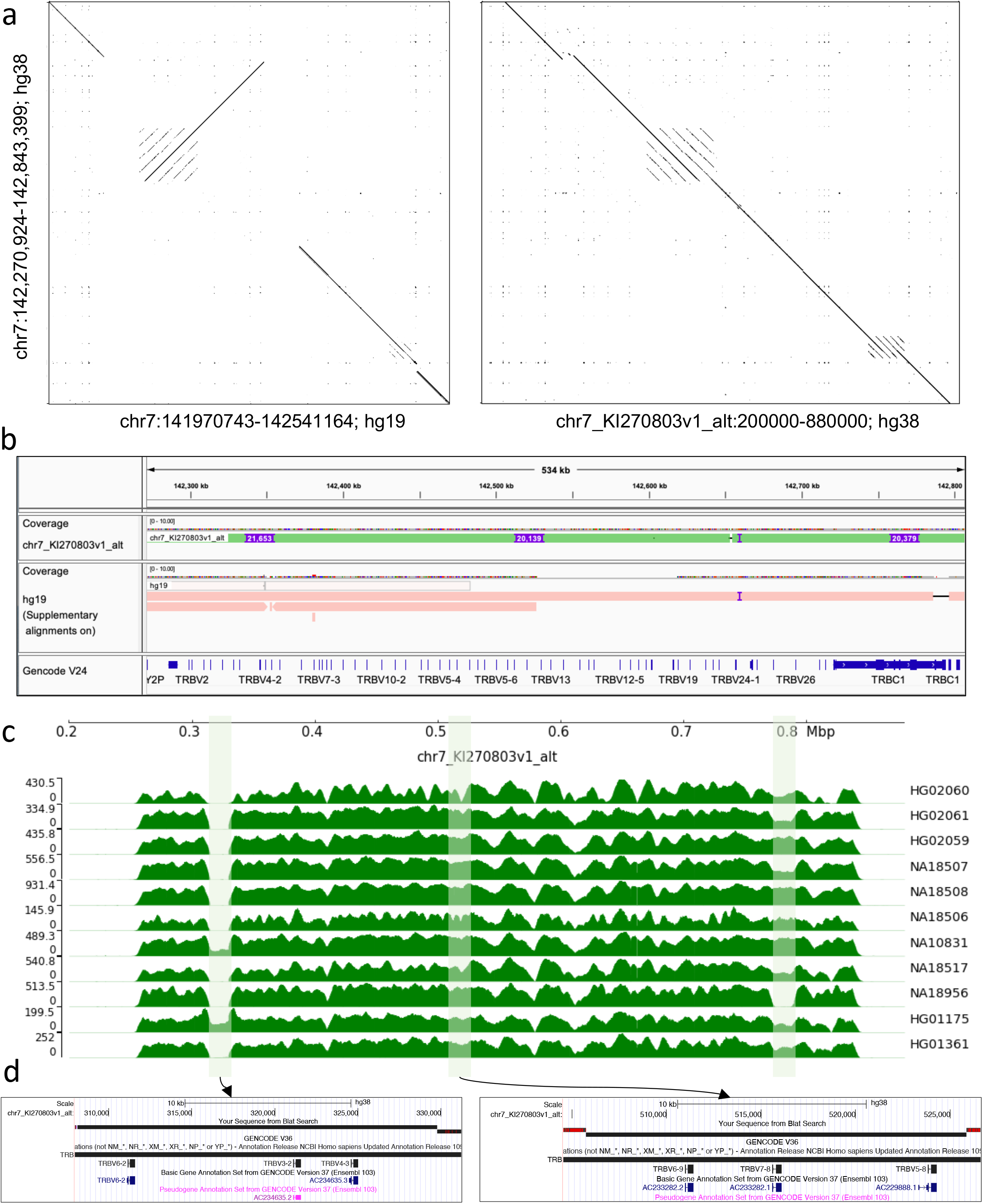
Comparison of TRB haplotypes in GRCh38/hg38 and GRGRh37/hg19. (**A**) Dot plots and (**B**) alignments between TRB loci GRCh38/hg38 chromosome 7 and GRCh37/hg19 chromosome 7 and GRCh38/hg38 chr7_KI270803v1_alt. (**C**) HiFi read coverage in all samples for the chr7_KI270803v1_alt assembly. Shaded regions indicate the three ~20 Kb insertions in chr7_KI270803v1_alt. (**D**) UCSC genome browser screenshots of chr7_KI270803v1_alt showing genes within the insertions.

### Detection of TRA/D and TRB alleles

A critical step in analyzing TCR repertoire sequencing data is the alignment of reads to a TCR germline gene database in order to identify the V, D and J allele present in a given read. Therefore, it is important to utilize a complete and accurate allele database(9). To determine the extent to which targeted long-read capture sequencing can help complete TCR germline databases, we first genotyped the alleles in both trios to measure the genotyping accuracy and then genotyped the remaining samples (Figure 5; Supplementary Table 1). All (n=207) TRA V and J, and TRB V, D and J alleles identified in HG02059 were correctly identified in the parents. The same was true for NA18506, except for the geneTRAJ18; in this case an additional allele was incorrectly identified in the assembly. However, direct use of the mapped HiFi reads facilitated characterization of the correct TRAJ18 allele.

**Figure 5.**
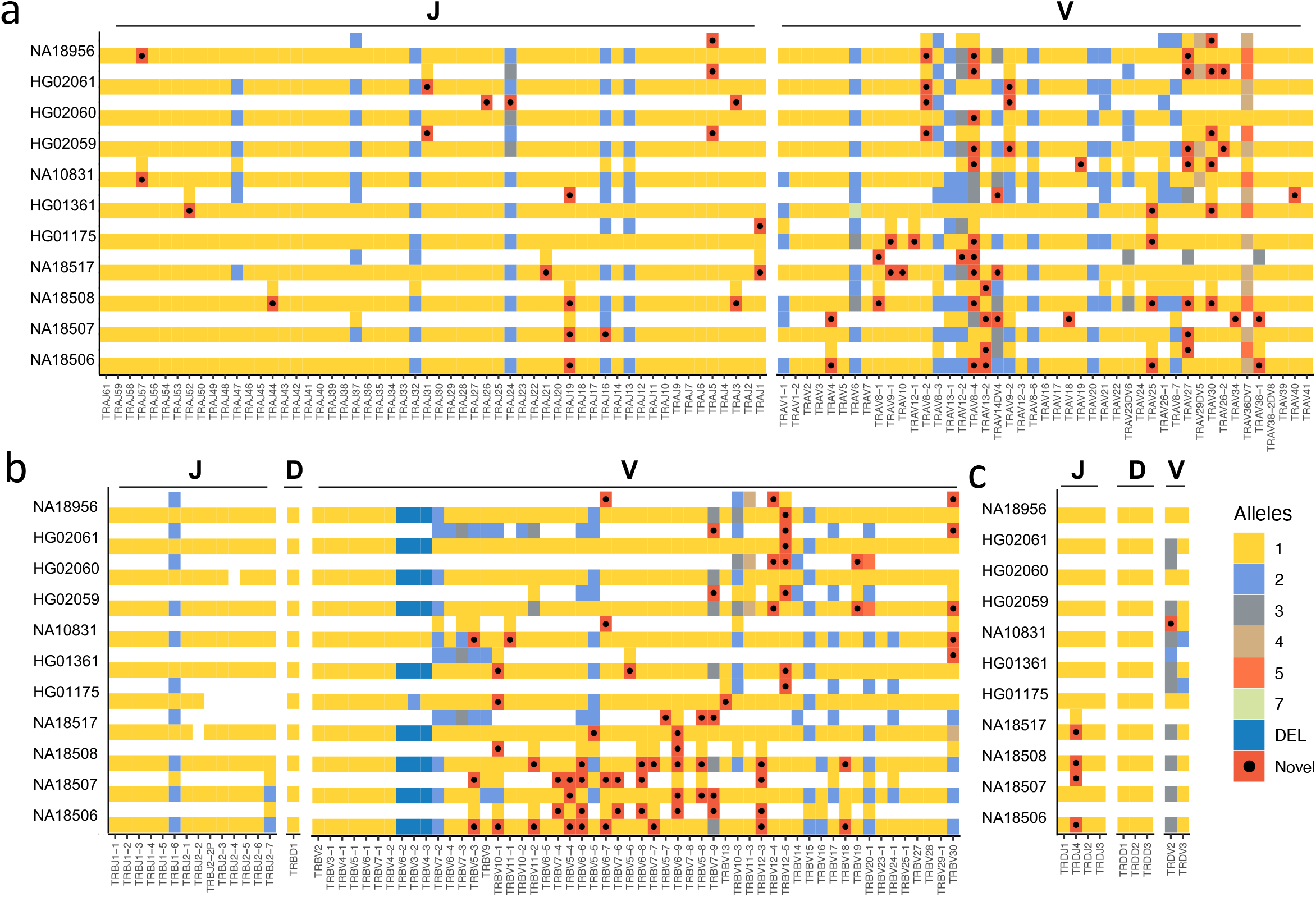
TRA/D and TRB allelic diversity. Gene alleles resolved from (**A**) TRA/D and (**B**) TRB long-read assemblies. Each color represents an allele or a deleted gene allele. Alleles marked by black dots are those not documented in IMGT. Genes with two distinct alleles are indicated by the presence of two filled boxes.

We next genotyped alleles across all nine unrelated samples (Figure 5; Supplementary Table 2). The average number of alleles observed per TRA V and J gene was 1.24 and 1.06, respectively, 1.19, 1 and 1.06 for TRB V, D and J genes, and 1.33, 1 and 1.06 for TRD V, D and J genes. This equated to an average of 26.7 heterozygous genes per sample. Notably, however, patterns of homozygosity and heterozygosity were biased toward particular genes in the TRA/D and TRB loci. For almost half of the genes (88 out 179) among TRA/D and TRB, no allelic variants were observed across all 9 samples. In contrast, for 7 genes (*TRAV27, TRAV36DV7, TRBJ1-6, TRBV10-3, TRBV30, TRAV12-2* and *TRAV8-4*), we observed heterozygous allele calls in at least six individuals.

We next evaluated the occurrence of novel alleles among our samples (*i.e.*, alleles not found in IMGT; (Supplementary Table 3). In TRA, TRB and TRD, there were 42, 35 and 2 novel alleles, respectively. The gene with the most novel alleles was TRAV8-4 (n=5). Across all the novel alleles, 13 (16%) were found in the assemblies and supported by >10 HiFi reads in 2 or more individuals. Thus, 66 of the putative novel alleles here were only found in one of the 9 unrelated samples; nonetheless, in all cases, these alleles were supported by both assemblies and >10 HiFi reads in each sample dataset. Given our estimates of assembly accuracy, and TR gene/allele genotyping recall in the trio datasets, we suspect these 66 alleles are likely genuine. Providing additional support, 7 of the 35 TRB alleles were identical to those recently reported in AIRR-seq germline inference study(9). Taken together, the 79 putative novel alleles identified here have the potential to increase the alleles available in IMGT by 23%, from a total of 341 to 420 functional/open-reading frame alleles. While all samples contain novel alleles, samples with African ancestry contain the most novel alleles empirically (Figure 5). This is likely due to both more genetic variation found within African populations and the underrepresentation of African samples in immunogenomic databases(20, 26).

Many recent studies have released TR allele databases with alleles derived from 1KG VCF files. One such database is the pmTRIG(27). Caution surrounding the use of short-read datasets for curating allelic variation in the immune receptor gene loci has been raised(28, 29). We checked to determine how many of our novel alleles were present in pmTRIG. Critically, the pmTRIG database used 2,548 1KG samples, including 9 samples in this study. However, we found that only 32 (76%), 16 (46%) and 0 (0%) of the novel TRA, TRB and TRD alleles, respectively, identified in the current study were present in pmTRIG. Consistent with our comparison of short- and long-read variant callsets (Figure 3d,e), this demonstrated that short-read datasets derived with next generation sequencing technologies might not be capable of detecting all novel alleles in the TRA/D and TRB regions, and likely results in erroneous allelic variant calls.

## Discussion

TCRs are critical to T cell function and the adaptive immune response. Although several studies have uncovered evidence that genetics plays a role in shaping the T cell repertoire, only two GWAS have implicated the TCR loci in disease pathogenesis(10–12). In addition, despite the importance of TCR repertoire sequencing and profiling in disease research, genetic variation in the TR loci has not been extensively characterized, including understanding the extent and types of variation within TR V, D, and J gene segments (9, 22, 30). We therefore extended our published immunogenomics framework to selectively sequence and assemble the human TR loci using long-read sequencing in two trios and five unrelated individuals spanning African, Asian, South American and European populations. Collectively, this dataset significantly expands the diversity and number of available annotated long-read haplotype assemblies for the TRA/D and TRB loci.

First, our analysis showed that the TCR loci can be effectively captured, sequenced and assembled into phased diploid assemblies using long-read data. Importantly, across all samples, >99.9% of bases across both TRA/D and TRB loci were spanned by HiFi reads, and use of read-based variant phasing allowed us to generate haploid-specific assembly contigs, in some cases extending up to 653 Kb in length. Critically, in trio probands fully-phased assemblies were possible, indicating that when phased variants are available, our method allows for the complete reconstruction of haplotype assemblies spanning these loci. Consistent with our previous analysis of the human IGH locus using this approach(17), comparison of assemblies from probands and parents of two trios demonstrated assembly per base accuracies >99.9%. This assembly accuracy was reflected in variant call sets between parental and proband samples as well, in which >99% of SNP genotypes followed Mendelian inheritance patterns. Together, these initial analyses in trios highlighted the utility of using our approach for deeper characterization of TRA/D and TRB genetic diversity in an extended sample set.

Focusing on the analysis of 9 unrelated samples of diverse ancestries, we showed that long-read assemblies could be used to comprehensively detect SNPs, indels and SVs. Similar to observations made in the IG loci(17, 31), although perhaps less extensive, we noted that genetic diversity in the TR loci was elevated, represented by significantly higher levels of SNP density relative to the chromosome 7 and 14 averages. Interestingly, densities of SNPs were slightly higher in TRA/D relative to TRB, but in contrast, more SVs were detected in the TRB locus. However, it is notable that the number of polymorphic gene-containing SVs in the TR loci collectively is fewer than currently described in the IG loci, particularly IGH. This is in line with previous suggestions that sequence evolution within the TR loci has been influenced less by SVs(32). Importantly, one of the gene deletions in TRB was very common in the samples studied here, present in a homozygous state in all 9 unrelated individuals sequenced. Being able to extend our approach to a greater number of individuals will be useful for more fully resolving SVs involving TCR genes, and assessing their frequencies across human populations. This will likely be the case with SNP detection and genotyping as well. Our analysis revealed that a large proportion of SNPs identified in our samples lacked allele frequency data in dbSNP. Many of these variants were difficult to genotype accurately using older short-read datasets from the 1KGP. While the utilization of higher coverage and quality short-read datasets performed better, our analysis indicated that the informed use of reference assembly files that account for particular SVs and alternate haplotypes is critical to ensure read mapping and genotyping accuracy. Future work should include the reanalysis of sequencing data utilizing a proper reference, or potentially a reference graph genome capable of integrating different haplotypes(33–35). A potential path forward includes a large-scale population long-read sequencing analysis of TR loci to generate completely phased haplotypes that will serve as a framework for detection and imputation of variants using previously generated sequencing datasets(34).

Our characterization of TR haplotypes extended to annotation of TR genes across donors. Having complete haplotypes facilitated the discovery of extensive allelic variation, including the presence of many novel TRA/D and TRB alleles. The fact that we identified 79 undocumented (non-IMGT) alleles in only 9 unrelated individuals highlights the severe deficits that currently exist in germline databases. Considering our findings alongside other recent efforts underlines the work that remains to be done to fully catalog TR gene diversity in the human population(9, 26). At present, the impact that these missing germline genes and alleles has on the analysis and interpretation of TCR repertoire sequencing studies is not clear, but would be expected to have profound effects on germline gene/allele assignment efforts, similar to what has been noted for IG repertoires. As observed in the IG loci(36–39), it will be interesting to understand whether the inter-individual diversity uncovered in the human TR loci will extend to non-human species as well.

Ultimately, we argue that improving the characterization of genetic diversity in the TR loci will shed light on the role of these regions in driving key functions of TCRs and T cells in a variety of disease and clinical contexts. This study demonstrates the effectiveness of our approach over other existing methods. As we have shown for the IG loci, our approach is scalable, offering the opportunity to utilize it in high-throughput fashion to sequence 100’s to 1000’s of samples. This would allow for the discovery and characterization of diversity in both coding and non-coding (*e.g.*, regulatory) regions of these critical immune loci at an unprecedented scale, with the potential to more fully catalog TR variants, akin to what has been done for the MHC/HLA genes. Haplotyping and genotyping of the MHC/HLA loci is standard practice in many immune studies, thus, our method provides an opportunity to similarly operationalize genotyping of TR genes, including efforts to do this in conjunction with MHC/HLA typing to better understand genetic impacts on the function of TCR-MHC interactions. Additionally, partnering our method with adaptive immune receptor repertoire (AIRR) sequencing can facilitate the identification of TR variants that impact the composition of the TCR repertoire(9, 40), as well as cross-talk between B and T cells, e.g. T cell dependent B cell activation. We expect such studies to become more common as the use of AIRR sequencing becomes more pervasive in the research and clinical arenas.

## References

1. Kumar, B.V., Connors, T.J. and Farber, D.L. (2018) Human T Cell Development, Localization, and Function throughout Life. Immunity, 48, 202–213.

2. The T Cell Receptor FactsBook (2001) 10.1016/b978-0-12-441352-8.x5000-0.

3. Roth, D.B. (2015) V(D)J Recombination: Mechanism, Errors, and Fidelity. Mobile DNA III, 10.1128/9781555819217.ch14.

4. Nikolich-Žugich, J., Slifka, M.K. and Messaoudi, I. (2004) The many important facets of T-cell repertoire diversity. Nat. Rev. Immunol., 4, 123–132.

5. Zvyagin, I.V., Pogorelyy, M.V., Ivanova, M.E., Komech, E.A., Shugay, M., Bolotin, D.A., Shelenkov, A.A., Kurnosov, A.A., Staroverov, D.B., Chudakov, D.M., et al. (2014) Distinctive properties of identical twins’ TCRrepertoires revealed by high-throughput sequencing. Proc. Natl. Acad. Sci. U. S. A., 111, 5980–5985.

6. Sharon, E., Sibener, L.V., Battle, A., Fraser, H.B., Christopher Garcia, K. and Pritchard, J.K. (2016) Genetic variation in MHC proteins is associated with T cell receptor expression biases. Nature Genetics, 48, 995–1002.

7. Klein, L., Kyewski, B., Allen, P.M. and Hogquist, K.A. (2014) Positive and negative selection of the T cell repertoire: what thymocytes see (and don’t see). Nature Reviews Immunology, 14, 377–391.

8. Gras, S., Chen, Z., Miles, J.J., Liu, Y.C., Bell, M.J., Sullivan, L.C., Kjer-Nielsen, L., Brennan, R.M., Burrows, J.M., Neller, M.A., et al. (2010) Allelic polymorphism in the T cell receptor and its impact on immune responses. Journal of Experimental Medicine, 207, 1555–1567.

9. Omer, A., Peres, A., Rodriguez, O.L., Watson, C.T., Lees, W., Polak, P., Collins, A.M. and Yaari, G. (2022) T cell receptor beta germline variability is revealed by inference from repertoire data. Genome Med., 14, 2.

10. Hallmayer, J., Faraco, J., Lin, L., Hesselson, S., Winkelmann, J., Kawashima, M., Mayer, G., Plazzi, G., Nevsimalova, S., Bourgin, P., et al. (2009) Narcolepsy is strongly associated with the T-cell receptor alpha locus. Nat. Genet., 41, 708–711.

11. Han, F., Faraco, J., Dong, X.S., Ollila, H.M., Lin, L., Li, J., An, P., Wang, S., Jiang, K.W., Gao, Z.C., et al. (2013) Genome wide analysis of narcolepsy in China implicates novel immune loci and reveals changes in association prior to versus after the 2009 H1N1 influenza pandemic. PLoS Genet., 9, e1003880.

12. O’Brien, R.P., Phelan, P.J., Conroy, J., O’Kelly, P., Green, A., Keogan, M., O’Neill, D., Jennings, S., Traynor, C., Casey, J., et al. (2013) A genome-wide association study of recipient genotype and medium-term kidney allograft function.Clin. Transplant., 27, 379–387.

13. Matsuda, F., Ishii, K., Bourvagnet, P., Kuma, K. i., Hayashida, H., Miyata, T. and Honjo, T. (1998) The complete nucleotide sequence of the human immunoglobulin heavy chain variable region locus. J. Exp. Med., 188, 2151–2162.

14. Rowen, L., Koop, B.F. and Hood, L. (1996) The Complete 685-Kilobase DNA Sequence of the Human beta T Cell Receptor Locus. Science, 272, 1755–1762.

15. Watson, C.T. and Breden, F. (2012) The immunoglobulin heavy chain locus: genetic variation, missing data, and implications for human disease. Genes Immun., 13, 363–373.

16. Zhao, X., Collins, R.L., Lee, W.-P., Weber, A.M., Jun, Y., Zhu, Q., Weisburd, B., Huang, Y., Audano, P.A., Wang, H., et al. (2021) Expectations and blind spots for structural variation detection from long-read assemblies and short-read genome sequencing technologies. Am. J. Hum. Genet., 108, 919–928.

17. Rodriguez, O.L., Gibson, W.S., Parks, T., Emery, M., Powell, J., Strahl, M., Deikus, G., Auckland, K., Eichler, E.E., Marasco, W.A., et al. (2020) A Novel Framework for Characterizing Genomic Haplotype Diversity in the Human Immunoglobulin Heavy Chain Locus. Front. Immunol., 11, 2136.

18. Kong, A., Gudbjartsson, D.F., Sainz, J., Jonsdottir, G.M., Gudjonsson, S.A., Richardsson, B., Sigurdardottir, S., Barnard, J., Hallbeck, B., Masson, G., et al. (2002) A high-resolution recombination map of the human genome. Nature Genetics, 31, 241–247.

19. Gibson, J., Morton, N.E. and Collins, A. (2006) Extended tracts of homozygosity in outbred human populations. Hum. Mol. Genet., 15, 789–795.

20. 1000 Genomes Project Consortium, Auton, A., Brooks, L.D., Durbin, R.M., Garrison, E.P., Kang, H.M., Korbel, J.O., Marchini, J.L., McCarthy, S., McVean, G.A., et al. (2015) A global reference for human genetic variation. Nature, 526, 68–74.

21. Byrska-Bishop, M., Evani, U.S., Zhao, X., Basile, A.O., Abel, H.J., Regier, A.A., Corvelo, A., Clarke, W.E., Musunuri, R., Nagulapalli, K., et al. High coverage whole genome sequencing of the expanded 1000 Genomes Project cohort including 602 trios. 10.1101/2021.02.06.430068.

22. Zhang, J.-Y., Roberts, H., Flores, D.S.C., Cutler, A.J., Brown, A.C., Whalley, J.P., Mielczarek, O., Buck, D., Lockstone, H., Xella, B., et al. (2021) Using de novo assembly to identify structural variation of eight complex immune system gene regions. PLoS Comput. Biol., 17, e1009254.

23. Lin, M.-J., Lin, Y.-C., Chen, N.-C., Luo, A.C., Lai, S.-K., Hsu, C.-L., Hsu, J.S., Chen, C.-Y., Yang, W.-S. and Chen, P.-L. Profiling Germline Adaptive Immune Receptor Repertoire with gAIRR Suite. 10.1101/2020.11.27.399857.

24. Steinberg, K.M., Schneider, V.A., Graves-Lindsay, T.A., Fulton, R.S., Agarwala, R., Huddleston, J., Shiryev, S.A., Morgulis, A., Surti, U., Warren, W.C., et al. (2014) Single haplotype assembly of the human genome from a hydatidiform mole. Genome Res., 24, 2066–2076.

25. Chaisson, M.J.P., Huddleston, J., Dennis, M.Y., Sudmant, P.H., Malig, M., Hormozdiari, F., Antonacci, F., Surti, U., Sandstrom, R., Boitano, M., et al. (2015) Resolving the complexity of the human genome using single-molecule sequencing. Nature, 517, 608–611.

26. Peng, K., Safonova, Y., Shugay, M., Popejoy, A.B., Rodriguez, O.L., Breden, F., Brodin, P., Burkhardt, A.M., Bustamante, C., Cao-Lormeau, V.-M., et al. (2021) Diversity in immunogenomics: the value and the challenge. Nat. Methods, 18, 588–591.

27. Khatri, I., Berkowska, M.A., van den Akker, E.B., Teodosio, C., Reinders, M.J.T. and van Dongen, J.J.M. Population matched (PM) germline allelic variants of immunoglobulin (IG) loci: New pmIG database to better understand*IG*repertoire and selection processes in disease and vaccination. 10.1101/2020.04.09.033530.

28. Watson, C.T., Matsen, F.A., Jackson, K.J.L., Bashir, A., Smith, M.L., Glanville, J., Breden, F., Kleinstein, S.H., Collins, A.M. and Busse, C.E. (2017) Comment on ‘A Database of Human Immune Receptor Alleles Recovered fromPopulation Sequencing Data’. J. Immunol., 198.

29. Collins, A.M., Peres, A., Corcoran, M.M., Watson, C.T., Yaari, G., Lees, W.D. and Ohlin, M. (2021) Commentary on Population matched (pm) germline allelic variants of immunoglobulin (IG) loci: relevance in infectious diseases and vaccination studies in human populations. Genes Immun., 22.

30. Mackelprang, R., Livingston, R.J., Eberle, M.A., Carlson, C.S., Yi, Q., Akey, J.M. and Nickerson, D.A. (2006) Sequence diversity, natural selection and linkage disequilibrium in the human T cell receptor alpha/delta locus. Hum. Genet., 119.

31. Watson, C.T., Steinberg, K.M., Huddleston, J., Warren, R.L., Malig, M., Schein, J., Willsey, A.J., Joy, J.B., Scott, J.K., Graves, T.A., et al. (2013) Complete haplotype sequence of the human immunoglobulin heavy-chain variable, diversity, and joining genes and characterization of allelic and copy-number variation. Am. J. Hum. Genet., 92, 530–546.

32. Luo, S., Yu, J.A., Li, H. and Song, Y.S. (2019) Worldwide genetic variation of the IGHV and TRBV immune receptor gene families in humans. Life Sci Alliance, 2.

33. Paten, B., Novak, A.M., Eizenga, J.M. and Garrison, E. (2017) Genome graphs and the evolution of genome inference. Genome Res., 27, 665–676.

34. Ebler, J., Ebert, P., Clarke, W.E., Rausch, T., Audano, P.A., Houwaart, T., Mao, Y., Korbel, J.O., Eichler, E.E., Zody, M.C., et al. (2022) Pangenome-based genome inference allows efficient and accurate genotyping across a wide spectrum of variant classes. Nat. Genet., 54, 518–525.

35. Rakocevic, G., Semenyuk, V., Lee, W.-P., Spencer, J., Browning, J., Johnson, I.J., Arsenijevic, V., Nadj, J., Ghose, K., Suciu, M.C., et al. (2019) Fast and accurate genomic analyses using genome graphs. Nat. Genet., 51, 354–362.

36. Kaduk, M., Corcoran, M. and Karlsson Hedestam, G.B. (2022) Addressing IGHV Gene Structural Diversity Enhances Immunoglobulin Repertoire Analysis: Lessons From Rhesus Macaque. Front. Immunol., 13, 818440.

37. Watson, C.T., Kos, J.T., Gibson, W.S., Newman, L., Deikus, G., Busse, C.E., Smith, M.L., Jackson, K.J. and Collins, A.M. (2019) A comparison of immunoglobulin IGHV, IGHD and IGHJ genes in wild-derived and classical inbred mouse strains. Immunol. Cell Biol., 97, 888–901.

38. Cirelli, K.M., Carnathan, D.G., Nogal, B., Martin, J.T., Rodriguez, O.L., Upadhyay, A.A., Enemuo, C.A., Gebru, E.H., Choe, Y., Viviano, F., et al. (2020) Slow Delivery Immunization Enhances HIV Neutralizing Antibody and Germinal Center Responses via Modulation of Immunodominance. Cell, 180, 206.

39. Kos, J.T., Safonova, Y., Shields, K.M., Silver, C.A., Lees, W.D., Collins, A.M. and Watson, C.T. Characterization of Extensive Diversity In Immunoglobulin Light Chain Variable Germline Genes Across Biomedically Important Mouse Strains. 10.1101/2022.05.01.489089.

40. Russell, M.L., Souquette, A., Levine, D.M., Schattgen, S.A., Kaitlynn Allen, E., Kuan, G., Simon, N., Balmaseda, A., Gordon, A., Thomas, P.G., et al. (2022) Combining genotypes and T cell receptor distributions to infer genetic loci determining V(D)J recombination probabilities. eLife, 11.

41. Chaisson, M.J. and Tesler, G. (2012) Mapping single molecule sequencing reads using basic local alignment with successive refinement (BLASR): application and theory. BMC Bioinformatics, 13, 238.

42. Martin, M., Patterson, M., Garg, S., Fischer, S.O., Pisanti, N., Klau, G.W., Schöenhuth, A. and Marschall, T. WhatsHap: fast and accurate read-based phasing. 10.1101/085050.

43. Rodriguez, O.L., Ritz, A., Sharp, A.J. and Bashir, A. (2020) MsPAC: a tool for haplotype-phased structural variant detection. Bioinformatics, 36, 922–924.

44. Nurk, S., Walenz, B.P., Rhie, A., Vollger, M.R., Logsdon, G.A., Grothe, R., Miga, K.H., Eichler, E.E., Phillippy, A.M. and Koren, S. (2020) HiCanu: accurate assembly of segmental duplications, satellites, and allelic variants from high-fidelity long reads. Genome Res., 30, 1291–1305.

45. Lassmann, T. (2019) Kalign 3: multiple sequence alignment of large data sets. Bioinformatics, 10.1093/bioinformatics/btz795.

