## Supplementary Figures for "Targeted long-read sequencing facilitates phased diploid assembly and genotyping of the human T cell receptor alpha, delta and beta loci"

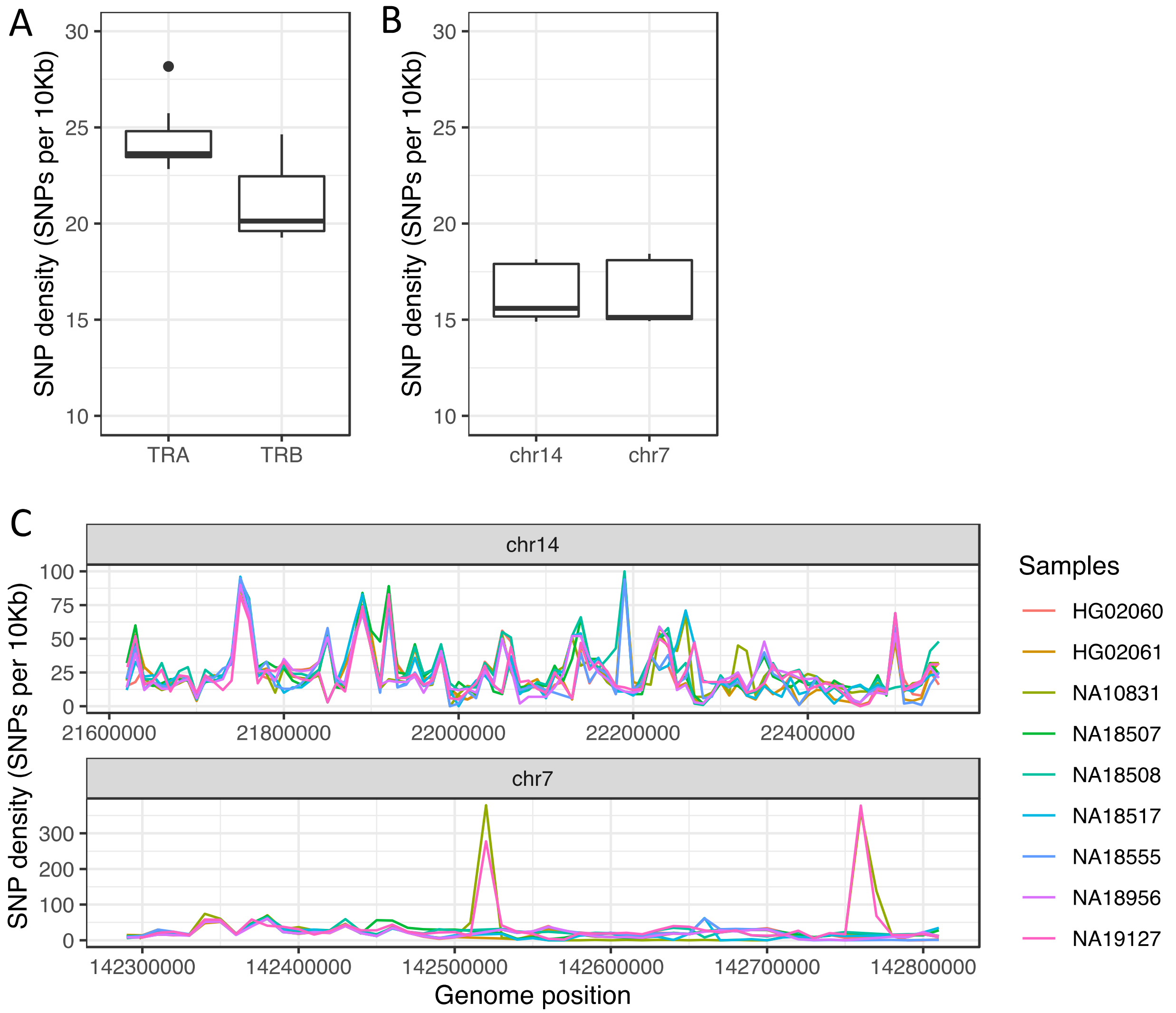


**Supplementary Figure 1. SNP density in TRA/D and TRB alleles.** SNP density in (**A**) TRA/D, TRB and (**B**) across chromosomes 14 and 7, excluding TR loci. **(C)** The number of SNPs per 10 Kb windows across TRA/D and TRB.

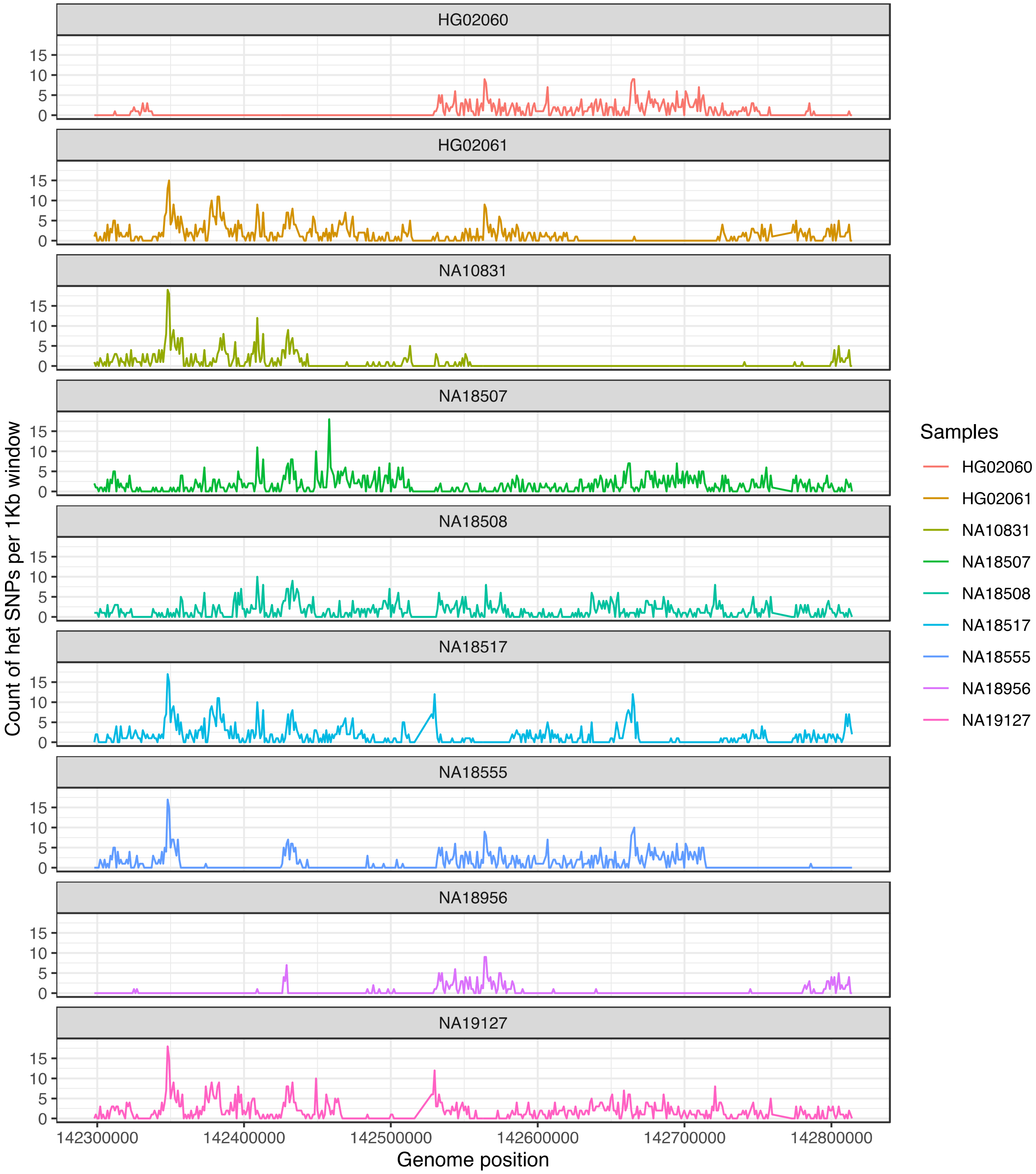


**Supplementary Figure 2. Number of heterozygous SNPs in TRB per 1Kb window.**

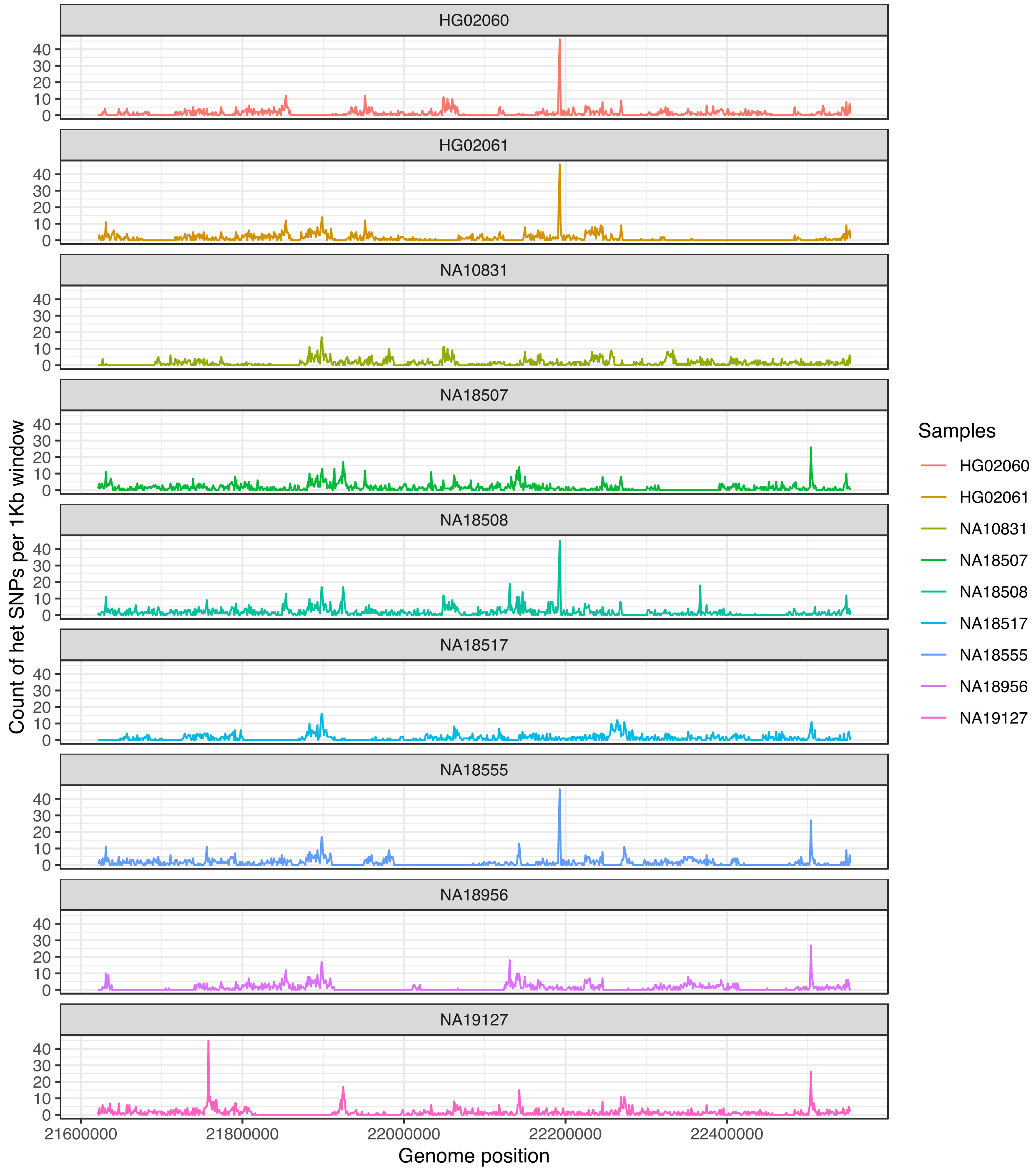


**Supplementary Figure 3. Number of heterozygous SNPs in TRA/D per 1Kb window.**
